# Design, construction and optimization of formaldehyde growth biosensors with broad application in Biotechnology

**DOI:** 10.1101/2023.06.29.547045

**Authors:** Karin Schann, Jenny Bakker, Maximilian Boinot, Pauline Kuschel, Hai He, Maren Nattermann, Tobias Erb, Arren Bar-Even, Sebastian Wenk

## Abstract

Formaldehyde is a key metabolite in natural and synthetic one-carbon metabolism as well as an important environmental toxin with high toxicity at low concentrations. To engineer efficient formaldehyde producing enzymes and to detect formaldehyde in industrial or environmental samples, it is important to establish highly sensitive, easy to use and affordable formaldehyde detection methods. Here, we transformed the workhorse bacterium *Escherichia coli* into biosensors that can detect a broad range of formaldehyde concentrations. Based on natural and promiscuous formaldehyde assimilation enzymes, we designed and engineered three different *E. coli* strains that depend on formaldehyde assimilation for cellular growth. After in depth characterization of these biosensors, we show that the formaldehyde sensitivity can be improved through adaptive laboratory evolution or modification of metabolic branch points. The metabolic engineering strategy presented in this work allowed the creation of *E. coli* biosensors that can detect formaldehyde in a concentration range from ∼30 μM to ∼13 mM. Using the most sensitive strain, we benchmarked the *in vivo* activities of different, widely used NAD-dependent methanol dehydrogenases, the rate-limiting enzyme in synthetic methylotrophy. We also show that the strains can grow upon external addition of formaldehyde indicating their potential use for applications beyond enzyme engineering. The formaldehyde biosensors developed in this study are fully genomic and can be used as plug and play devices for screening large enzyme libraries. Thus, they have the potential to greatly advance enzyme engineering and might even be used for environmental monitoring or analysis of industrial probes.

**Highlights:**

- Conversion of *E. coli* into three different formaldehyde growth biosensors
- Biosensors are fully genomic and grow robustly when formaldehyde is present
- Biosensors can detect formaldehyde concentrations ranging from ∼30 μM to ∼13 mM
- Benchmarking of biotechnological relevant methanol dehydrogenases reveals potential of biosensors for enzyme engineering
- Biosensors grow upon direct addition of formaldehyde indicating potential use in environmental or industrial settings

## Introduction

Formaldehyde is a simple, highly reactive molecule consisting of a central carbon atom bound by one oxygen and two hydrogen atoms (see **Figure 1**). Due to its small size, high solubility and volatility (gaseous under atmospheric conditions), formaldehyde spreads easily and enters the cell by diffusion through the cell membrane (Salthammer, 2022). Inside the cell, its high reactivity can lead to crosslinking of DNA and proteins, triggering mutagenesis and cell death (Bernardini et al., 2020; Reingruber and Pontel, 2018). Despite its high toxicity and carcinogenicity (Bosetti et al., 2008), formaldehyde remains indispensable for numerous industrial processes, including the production of textiles and various building materials (Reingruber and Pontel, 2018). Therefore, formaldehyde remains a serious hazard to the environment and human health.

**Figure 1:**
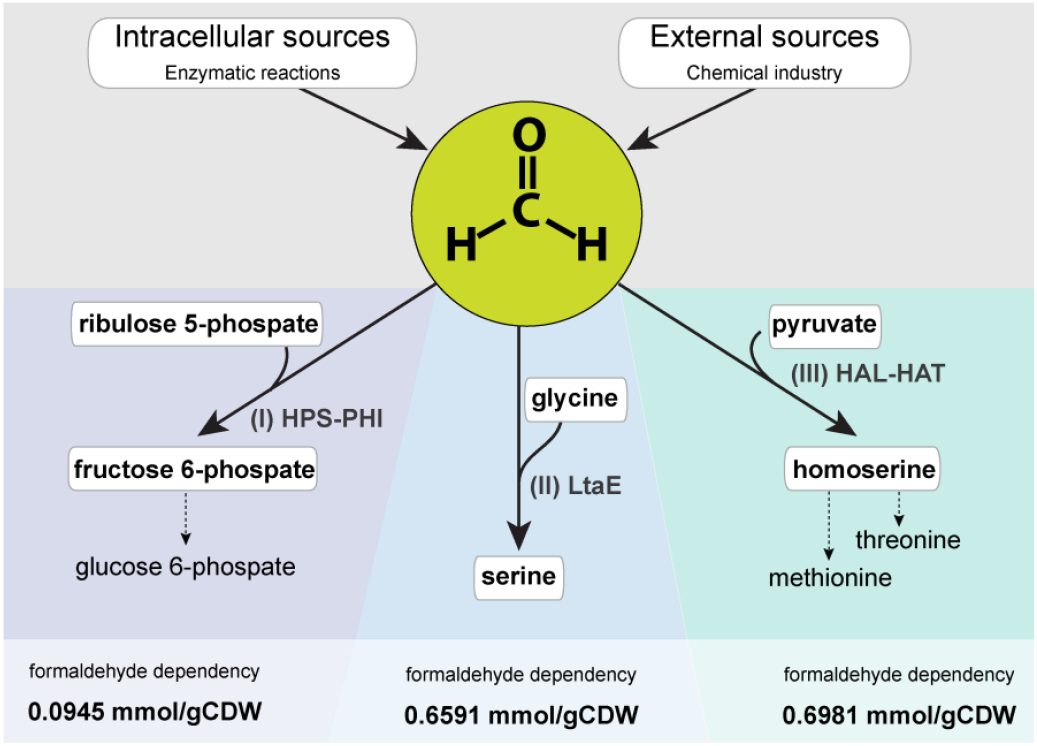
Formaldehyde detection via three different *E. coli* growth biosensors. Formaldehyde is a key intermediate in methylotrophic C_1_ metabolism and an important but highly toxic industrial chemical. Based on three formaldehyde assimilating enzymes, we designed, constructed and characterized *E. coli* formaldehyde growth biosensors that depend on formaldehyde assimilation via these reactions for cell growth. (I) The first biosensor is based on the RuMP cycle enzymes 3-hexulose-6-phosphate synthase (HPS) and 6-phospho-3-hexuloisomerase (PHI) to assimilate formaldehyde and ribulose 5-phosphate (Ru5P) into the essential metabolite fructose-6-phosphate (F6P). As F6P is converted to glucose 6-phosphate (G6P) the strain depends on formaldehyde assimilation for the generation of ∼4 % of its biomass (Frederick C. Neidhardt, John L. Ingraham, 1992). (II) The second formaldehyde biosensor is based on the promiscuous serine aldolase activity of *E. coli’s* low-specificity L-threonine aldolase (LtaE) to condense formaldehyde and glycine into the essential amino acid serine. Thus, the strain depends on formaldehyde assimilation for the generation of ∼2.1 % of its biomass (Frederick C. Neidhardt, John L. Ingraham, 1992). (III) The thirds biosensor is based on the promiscuous 4-hydroxy-2-oxobutanoate (HOB) aldolase (HAL) reaction of 2-keto-3-deoxy-L-rhamnonate aldolase (RhmA) followed by a HOB transaminase (HAT) reaction catalyzed by an aminotransferase which assimilate formaldehyde and pyruvate into homoserine. As homoserine is the precursor of the essential amino acids threonine and methionine, the strain depends on formaldehyde assimilation for the generation of ∼4.7 % of its biomass (Frederick C. Neidhardt, John L. Ingraham, 1992). The formaldehyde dependency of each strain (mmol/gCDW) was calculated by flux balance analysis based on the *E. coli* genome scale metabolic model and is indicated in the bottom of the figure.

On the other hand, formaldehyde is a central metabolite in methylotrophic one-carbon (C_1_) metabolism where methanol-derived formaldehyde is directly assimilated into biomass via the ribulose monophosphate (RuMP) cycle, the xylulose monophosphate (XuMP) cycle or the serine cycle (Antoniewicz, 2019; Zhang et al., 2017). As methanol can be produced electrochemically from CO_2_ (Bowker, 2019), it is expected to play a major role in a sustainable C_1_ bio-economy where native or synthetic methylotrophs convert methanol into value-added products (Claassens et al., 2019; Whitaker et al., 2015). Since the engineering of native methylotrophs for bioproduction can be challenging (Schada Von Borzyskowski et al., 2015), recent efforts have focused on engineering synthetic methylotrophy in biotechnological hosts like *Sacharomyces cerevisiae* and *Escherichia coli* (Chen et al., 2020; Espinosa et al., 2020; He et al., 2020; Keller et al., 2022; Kim et al., 2020). While this approach enabled growth of *E. coli* on methanol via the reductive glycine pathway (rGlyP) and the RuMP cycle, only the RuMP strain showed decent growth on methanol (doubling time ∼2 hrs and max. OD_600_ ∼2) (Chen et al., 2020; Keller et al., 2022). In the meantime, many other engineering projects did not achieve growth on methanol alone (e.g. He et al. 2018, 2020; Meyer et al. 2018; Yu and Liao 2018). A major bottleneck in engineering growth on methanol is the lack of efficient methanol dehydrogenases (MDHs) that can be heterologously expressed in microbial hosts. The widely used NAD-dependent MDHs are easy to express but display poor activities, including low catalytic rates and *K*_*m*_ values in the millimolar range. On the other hand, PQQ-dependent MDHs are very efficient, but difficult to express (Krüsemann et al., n.d.; Le et al., 2021). Thus, engineering efficient MDHs for synthetic methanol assimilation is still an open challenge in Biotechnology.

Another attractive sustainable feedstock for a C_1_ bio-economy is formate. Like methanol, it can be efficiently produced from CO_2_ (Yishai et al., 2016). A synthetic route from formate to formaldehyde has been proposed by Arren Bar-Even. In the suggested two enzyme-cascade, formate is either activated to formyl-CoA or formyl phosphate, which are subsequently reduced to formaldehyde (Bar-Even, 2016). This route, which does not seem to exist in nature, would vastly increase the number of formate assimilation routes as it could be combined with the above-mentioned natural formaldehyde assimilation routes or synthetic routes like the homoserine cycle or the formolase pathway (He et al., 2020; Siegel et al., 2015). So far, two studies could show the activity of the formyl-CoA route (Hu et al., 2022; Wang et al., 2021) and another recent study showed activity of the formyl-phosphate route by engineering a new-to-nature formyl phosphate reductase (Nattermann et al., 2023). However, achieving bacterial growth via either route is still pending.

To establish and optimize growth via formaldehyde, it is necessary to screen for formaldehyde producing enzymes (either natural or engineered). Assessing the catalytic activity in large libraries however requires the development of suitable screening systems. Here, molecular biosensors could be a valuable tool to detect enzymatic formaldehyde production. As such, a highly sensitive plasmid-based formaldehyde biosensor has been developed by Woolston et al. This sensor produces a fluorescence signal upon intracellular formaldehyde production and could be used for gene library screenings using fluorescence activated cell sorting (FACS) or other fluorescence-based methods (Woolston et al., 2018). However, this approach can be challenging, especially for large libraries, as it requires the use of sophisticated FACS equipment. Furthermore, plasmid related problems like copy number variations and plasmid incompatibility could be an issue when using a plasmid-based biosensor. A simpler and more robust way of detecting intracellular formaldehyde (fewer signal variations and background signal) could be achieved by engineering fully genomic growth-coupled formaldehyde biosensors. In these biosensors, enzyme activity is linked to cellular growth (Aslan et al., 2020; Orsi et al., 2021; Wenk et al., 2020). By strategically interrupting metabolism through gene deletions and expression of formaldehyde assimilating enzymes, growth under specific conditions is exclusively rescued upon sufficient enzymatic formaldehyde production. Such strains can directly select for the few variants in a library whose activity is high enough to produce formaldehyde at sufficient amounts, enabling rapid screening of large gene libraries. In this study, we designed, developed, and optimized formaldehyde growth biosensors that can detect a broad range of formaldehyde concentrations. First, we screened the metabolic landscape of *E. coli* for suitable formaldehyde entry points (natural and synthetic) and determined three formaldehyde assimilation reactions as the basis for the biosensor design. Based on these reactions, we designed three distinct growth biosensors that depend on the presence of intracellular formaldehyde for cellular growth: (I), the RuMP biosensor is based on the RuMP cycle enzymes 3-hexulose-6-phosphate synthase (HPS) and 6-phospho-3-hexuloisomerase (PHI) to assimilate formaldehyde and ribulose 5-phosphate (Ru5P) into fructose-6-phosphate (F6P). (II), the LtaE biosensor is based on the promiscuous activity of *E. coli’s* low-specificity L-threonine aldolase (LtaE) to condense formaldehyde and glycine into serine. (III) The HOB biosensor is based on the promiscuous 4-hydroxy-2-oxobutanoate (HOB) aldolase (HAL) reaction of *E. coli’s* 2-keto-3-deoxy-L-rhamnonate aldolase (RhmA) followed by a HOB transaminase (HAT) reaction which assimilate formaldehyde and pyruvate into homoserine (**Figure 1**). Using flux balance analysis (FBA), we estimated the formaldehyde dependency of the different strains (**Figure S1**). Then, we engineered the strains, validated their formaldehyde dependency and determined a suitable expression level of the formaldehyde-assimilating enzyme. In an adaptive laboratory evolution (ALE) experiment, we optimized the sensitivity of the LtaE biosensor. Furthermore, we also showed that the formaldehyde dependency of the HOB biosensor can be modulated by adding threonine to the medium. In detailed characterization experiments, we determined the formaldehyde sensitivity of each biosensor and provide insights to potential applications of the biosensors: By testing several widely used MDHs in the HOB biosensor, we benchmarked their *in vivo* activities demonstrating that the biosensors could be highly useful for enzyme discovery or engineering. We also show that two of the biosensors can grow upon direct addition of formaldehyde indicating their potential for industrial or environmental applications.

In Summary, we constructed and characterized a set of highly robust formaldehyde growth biosensors that can detect a broad range of formaldehyde concentrations and demonstrated their applicability for enzyme engineering and direct formaldehyde detection.

## Results

In the following, we describe the construction and characterization of the different formaldehyde biosensors. In all characterization experiments, a sarcosine oxidase (SoxA) from *Bacillus sp. B-0618* (SoxA; UniProt: P40859) was used as a formaldehyde producing enzyme. SoxA efficiently cleaves non-cytotoxic sarcosine (N-methyl-glycine) into formaldehyde and glycine and has been shown to robustly produce formaldehyde in previous studies (He et al., 2020, 2018). While methanol is surely the more desirable substrate for a C_1_ bio-economy, we determined it to be unsuitable as a formaldehyde precursor for the characterization experiments as its evaporation causes strong experimental variations. Furthermore, the available NAD-dependent MDHs present less favorable enzyme characteristics compared to SoxA.

Before going into detail about the different strains, we want to put forward our definition of formaldehyde sensitivity, which we use to evaluate the sensor strains. We define formaldehyde sensitivity as follows:

*“The formaldehyde sensitivity of a biosensor is defined by the lowest concentration of formaldehyde at which the strain shows a measurable growth signal (Max. OD*_*600*_ *> 0*.*1) within the experimental time frame (< 200 hours). The sensitivity inversely correlates with the concentration of detectable formaldehyde, i*.*e. the lower the formaldehyde concentration that can be detected by the strain, the higher its formaldehyde sensitivity. We note that in the case of two strains growing at the same concentration of formaldehyde, the strain which grows earlier and/or to a higher OD*_*600*_ *is considered more sensitive*.

In theory, the formaldehyde sensitivity of the biosensor strains could be determined by FBA (as indicated in **Figure 1** and **Figure S1**). As strains require formaldehyde (assimilation) for growth, their sensitivities could be deduced from the minimal required amount of formaldehyde per unit cell dry weight (gCDW) increase. While the calculation assumes all other factors are sufficient and optimal, this is not necessarily the case in the cellular environment (because of the availability of co-factors, enzyme expression levels, or stability of intermediates etc.), making experimental validation strictly necessary.

### The RuMP biosensor can detect formaldehyde in the millimolar range

Based on the natural formaldehyde assimilating enzymes of the RuMP cycle (HPS and PHI), we designed and constructed the RuMP biosensor with a low formaldehyde dependency (0.0945 mmol/gCDW as estimated by FBA) (**Figure 2A**). Through the deletions of the genes encoding for fructose 1,6-bisphosphatases 1 and 2 (*fbp and glpX*) as well as transketolases 1 and 2 (*tktA* and *tktB*), the strain is auxotrophic for fructose 6-phosphate (F6P) and glucose 6-phosphate (G6P) when cultivated on a carbon source from lower metabolism (e.g. succinate). These auxotrophies could be relieved if formaldehyde is condensed with ribulose 5-phosphate (Ru5P) (derived from xylose provided in the medium) and assimilated into F6P via HPS and PHI. Previously, it was suggested that HPS is the rate limiting enzyme in the conversion of formaldehyde and Ru5P into F6P (Kato et al., 2006). In this study, the *K*_*M*_ of HPS for formaldehyde was determined to be in the range of ∼0.5 to ∼2 mM which might directly influence the formaldehyde sensitivity of the biosensor. To increase the strain’s sensitivity for formaldehyde, the genes encoding for *frmRAB* and *zwf* were deleted. *frmRAB* encodes for the endogenous formaldehyde detoxification system of *E. coli* which oxidizes formaldehyde to formate. Its deletion removes a formaldehyde sink and increases the formaldehyde availability for assimilation by HPS. On the other hand, *zwf* encodes for the glucose 6-phosphate dehydrogenase which converts G6P into glucono-1,5-lactone 6-phosphate. Its deletion prevents G6P depletion and is thus expected to increase the sensitivity of the strain (He et al., 2018).

**Figure 2:**
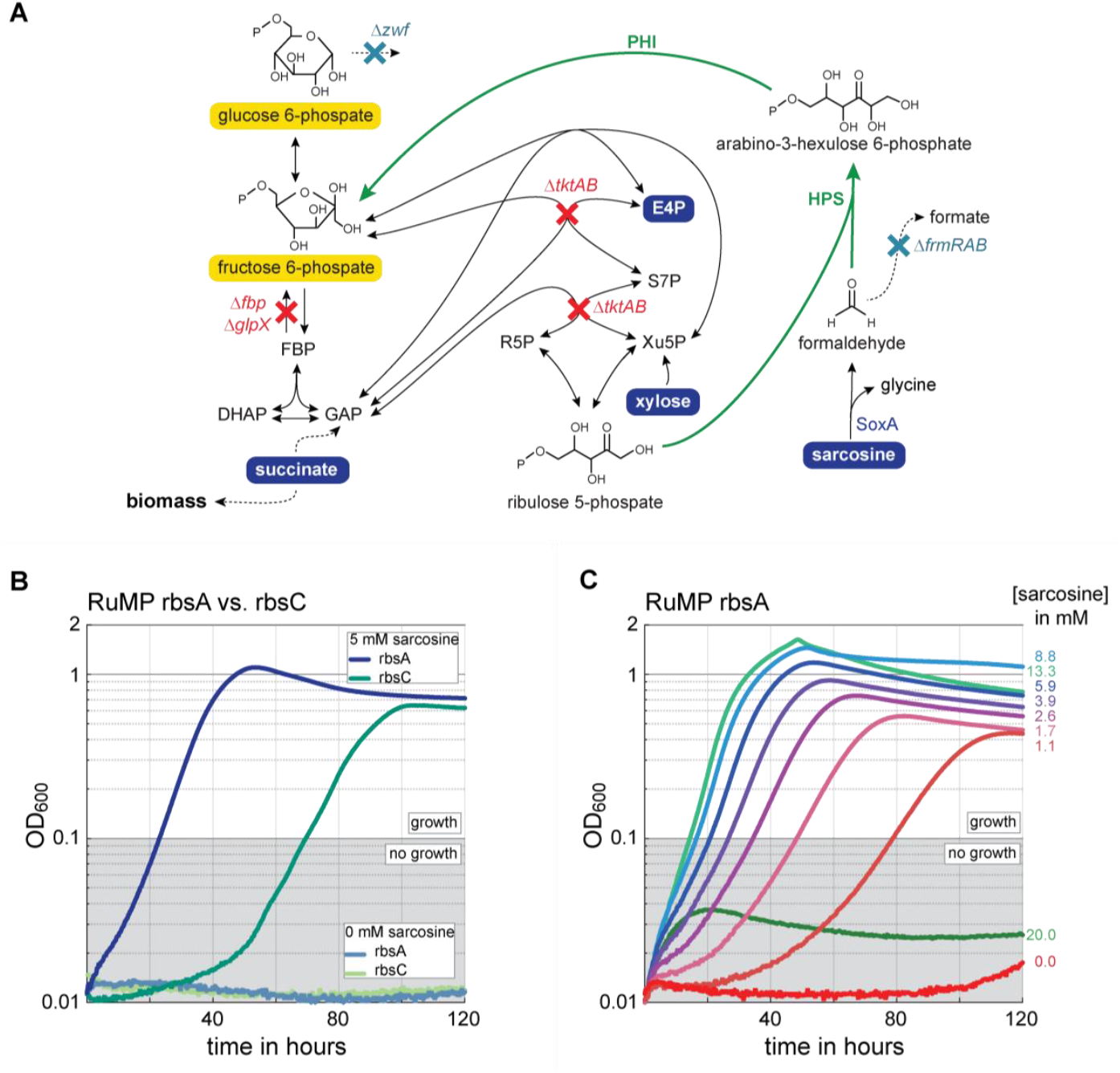
The RuMP biosensor can detect formaldehyde in the millimolar range. (A) Metabolic scheme of the RuMP biosensor. The strain uses the RuMP cycle enzymes HPS and PHI for the assimilation of formaldehyde and ribulose 5-phosphate (Ru5P) into fructose 6-phosphate (F6P). The strain was engineered to be auxotrophic for F6P and glucose 6-phosphate (G6P) through the indicated deletions (red crosses). To increase the sensitivity of the strain, potential sink reactions for formaldehyde and G6P were removed by deleting the formaldehyde detoxification system (*frmRAB*) and the G6P sink (*zwf*) (cyan crosses). To assimilate formaldehyde and Ru5P into F6P, HPS and PHI from *Bacillus methanolicus* were cloned into a synthetic operon and expressed from the genome using a strong constitutive promoter and either a medium or strong ribosome binding site (rbsC or rbsA, respectively). Additionally, SoxA was expressed from a plasmid to allow sarcosine conversion into formaldehyde. (B) The RuMP biosensor with either rbsA or rbsC was cultivated in minimal medium, supplemented with 19 mM succinate, 12 mM xylose, 2 mM glycine, E4P mix with and without the addition of 5 mM sarcosine. Only when sarcosine was added to the medium, the strains grew indicating their dependency on formaldehyde. (C) To characterize the RuMP biosensor in more detail, the strain with rbsA was cultivated with a sarcosine gradient ranging from 0 to 20 mM. The gradient revealed a correlation between growth and sarcosine concentration between 1.1 and 13.3 mM. Abbreviations: DHAP, dihydroxyacetone phosphate; E4P, erythrose 4-phosphate; FBP, fructose 1,6-bisphosphate; GAP, glyceraldehyde 3-phosphate; R5P, ribose 5-phosphate; S7P, sedoheptulose 7-phosphate; X5P, xylulose 5-phosphate; *fbp*, fructose 1,6-bisphosphatase 1; *glpX*, fructose 1,6-bisphosphatase 2; *tktAB*, transketolase 1 and 2; *frmRAB*, formaldehyde detoxification system; *zwf*, glucose 6-phosphate dehydrogenase; SoxA, sarcosine oxidase; HPS, 3-hexulose-6-phosphate synthase; PHI, 6-phospho-3-hexuloisomerase. Figure elements: yellow background: auxotrophy; blue circle: substrate; red cross: deletion causing auxotrophy; cyan cross: non-essential deletion; green arrow: formaldehyde assimilation flux; dashed arrow: multi enzymatic reaction.

After deletion of all the above-mentioned genes, the HPS and PHI genes were cloned into a synthetic operon either with a medium or a strong ribosome binding site (RBS) (rbsC and rbsA, respectively) and expressed from a safe spot in the genome (Bassalo et al., 2016) under the control of a strong constitutive promoter. To confirm the RuMP biosensor’s ability to detect intracellularly produced formaldehyde, we transformed the strains with a plasmid encoding a SoxA. Then, we cultivated both strains with a defined concentration of sarcosine to compare their formaldehyde sensitivity, which we derived from the growth phenotype. As the strain with the strong rbsA showed a higher formaldehyde sensitivity compared to the strain with the medium strength rbsC (**Figure 2B**), we used the RuMP biosensor with the rbsA for detailed characterization. Testing the strain with a sarcosine gradient revealed a formaldehyde detection range between 1 and 13 mM making the strain a suitable biosensor for high to medium formaldehyde concentrations (**Fig. 2C**).

### The LtaE biosensor can detect formaldehyde in the milli-to micromolar range

The design of the LtaE biosensor is based on the promiscuous serine aldolase activity of the *E. coli* enzyme LtaE (Contestabile et al., 2001). The enzyme naturally cleaves threonine into acetaldehyde and glycine but can also condense glycine and formaldehyde to serine *in vivo* (He et al., 2020). Expressing LtaE in a strain that cannot synthesize serine from 3-phosphoglycerate (Δ*serA*) or from glycine and methylene-THF (Δ*glyA*) should create a formaldehyde biosensor strain, that is only capable to grow on glucose and glycine when formaldehyde is present in the cell and can condense with glycine to form serine. Using FBA, we estimated that the LtaE biosensor has a higher formaldehyde dependency than the RuMP biosensor (0.6591 mmol/gCDW) suggesting it to be less sensitive to formaldehyde.

After deleting *serA, glyA* and *frmRAB* as well as the endogenous *ltaE* gene, we integrated *ltaE* with either a medium or a strong RBS (rbsC and rbsA, respectively) into a safe spot in the genome (Bassalo et al., 2016) and expressed it under the control of a strong constitutive promoter (**Figure 3A**). The resulting strains were transformed with a plasmid encoding for SoxA and tested for their ability to detect intracellularly produced formaldehyde. Both strains were cultivated with a defined concentration of sarcosine to compare their sensitivity. Also here, the strain expressing the formaldehyde assimilating enzyme with the strong rbsA showed a higher formaldehyde sensitivity compared to the rbsC strain (**Figure 3B**). Hence, subsequent experiments were carried out with the more sensitive LtaE biosensor with the rbsA.

**Figure 3:**
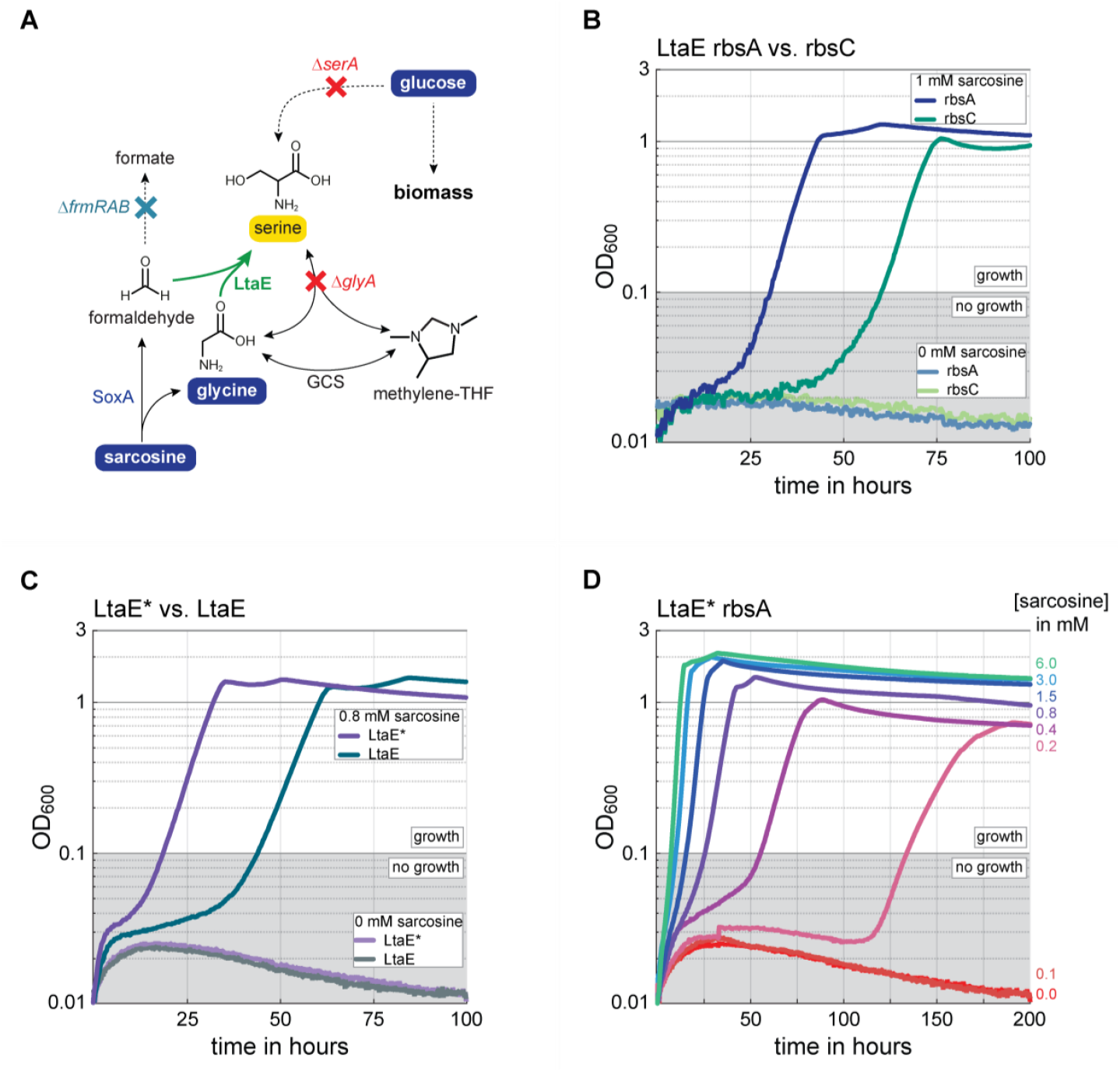
Enzyme evolution improves the LtaE biosensor and enables formaldehyde detection in the mili-to micromolar range. (A) Metabolic scheme of the LtaE biosensor. The strain is based on the promiscuous activity of the LtaE enzyme that condenses formaldehyde and glycine into serine. To make the strain dependent of formaldehyde assimilation, *serA* and *glyA* were deleted to create a serine auxotrophy. To increase the sensitivity of the strain, a formaldehyde sink was removed by deleting the endogenous formaldehyde detoxification system of *E. coli* (*frmRAB*). To assimilate formaldehyde and glycine into serine, LtaE from *E. coli* was expressed from the genome using a strong constitutive promoter and either a medium or strong ribosome binding site (rbsC or rbsA, respectively). SoxA was expressed from a plasmid to allow sarcosine conversion into formaldehyde. (B) The LtaE biosensor expressing *ltaE* either under the control of rbsA or rbsC was cultivated in minimal medium, supplemented with 10 mM glucose, 10 mM glycine, 50 μM MnCl_2_ with and without the addition of 1 mM sarcosine. (C) An amino acid exchange (C188Y) observed in an adaptive laboratory evolution experiment was introduced into LtaE creating the LtaE^*^ biosensor. Comparison of the LtaE and the LtaE^*^ biosensor with rbsA showed that the LtaE^*^ biosensor has a 2-fold faster growth when cultivated with 0.8 mM sarcosine. (D) For detailed characterization of the formaldehyde sensitivity, the LtaE^*^ biosensor was cultivated in a gradient of sarcosine concentrations ranging from 0 to 6 mM. The gradient revealed a correlation between growth and sarcosine concentration between 0.2 and 6 mM. Abbreviations: *glyA*, serine hydroxymethyl transferase; *serA*, phosphoglycerate dehydrogenase; *frmRAB*, formaldehyde detoxification system; SoxA, sarcosine oxidase; LtaE, threonine aldolase; GCV, glycine cleavage system. Figure elements: yellow background: auxotrophy; blue circle: substrate; red cross: deletion causing auxotrophy; cyan cross: non-essential deletion; green arrow: formaldehyde assimilation flux; dashed arrow: multi enzymatic reaction.

As the LtaE biosensor depends on the activity of only one enzyme to relieve its auxotrophy, we hypothesized that the strain could be “easily” improved by ALE. To this end, the LtaE biosensor was transformed with a MDH from *Corynebacterium glutamicum* (CgMDH) and continuously cultivated on glucose, glycine and decreasing concentrations of methanol in closed tubes. In this experiment, MDH was used instead of SoxA as the low turnover of MDH presumably yields lower intracellular formaldehyde levels, which may drive LtaE evolution towards a lower *K*_m_. The short-term tube evolution resulted in strains with improved methanol dependent growth, capable of growing with 50 mM methanol (**Figure S2**). When analyzing these strains via whole genome sequencing, we identified a point mutation in the *ltaE* gene that lead to a dedicated amino acid exchange in LtaE (C188Y). Although, the respective amino acid is not located near the active site of the enzyme, detailed *in vitro* characterization of both enzyme variants revealed a lowering in the *K*_m_ for glycine, yielding a two-fold improvement in *k*_cat_/*K*_m_ (**Figure S3**).

As the optimized LtaE biosensor carrying the C188Y mutation (from here on termed LtaE^*^ biosensor) showed a consistently higher formaldehyde sensitivity than the original LtaE biosensor (**Figure 3C**), we went on to characterize this strain in more detail. To this end, we conducted a growth experiment with a sarcosine gradient. The experiment revealed a formaldehyde detection range of the LtaE^*^ biosensor between 0.2 and 6 mM making the strain more sensitive than the RuMP biosensor and a suitable biosensor for medium to low formaldehyde concentrations (**Figure 3D**).

### The HOB biosensor can detect formaldehyde in the low micromolar range

The design of the HOB biosensor is based on the combined HAL and HAT reactions of the homoserine cycle (He et al., 2020). In the HAL reaction, pyruvate is condensed with formaldehyde to generate HOB, which is subsequently aminated to homoserine via the HAT reaction. In *E. coli*, the HAL reaction is promiscuously catalyzed by 2-keto-3-deoxy-L-rhamnonate aldolase (RhmA) (Hernandez et al., 2017), whereas the HAT reaction is catalyzed by numerous aminotransferases (Hernandez et al., 2017; Walther et al., 2018; Zhong et al., 2019). To create a formaldehyde biosensor based on the combined HAL and HAT, we disrupted the native homoserine biosynthesis pathway of *E. coli* (*Δasd*), thereby generating a threonine and methionine auxotrophic strain (**Figure 4A**). As the strain depends on 0.6981 mmol formaldehyde for the generation of one gCDW (as estimated by FBA), this strain should in theory be the least sensitive formaldehyde biosensor. However, as methionine and threonine cannot be interconverted by *E. coli*, supplementation of either amino acid could reduce the biomass dependency of the strain.

**Figure 4:**
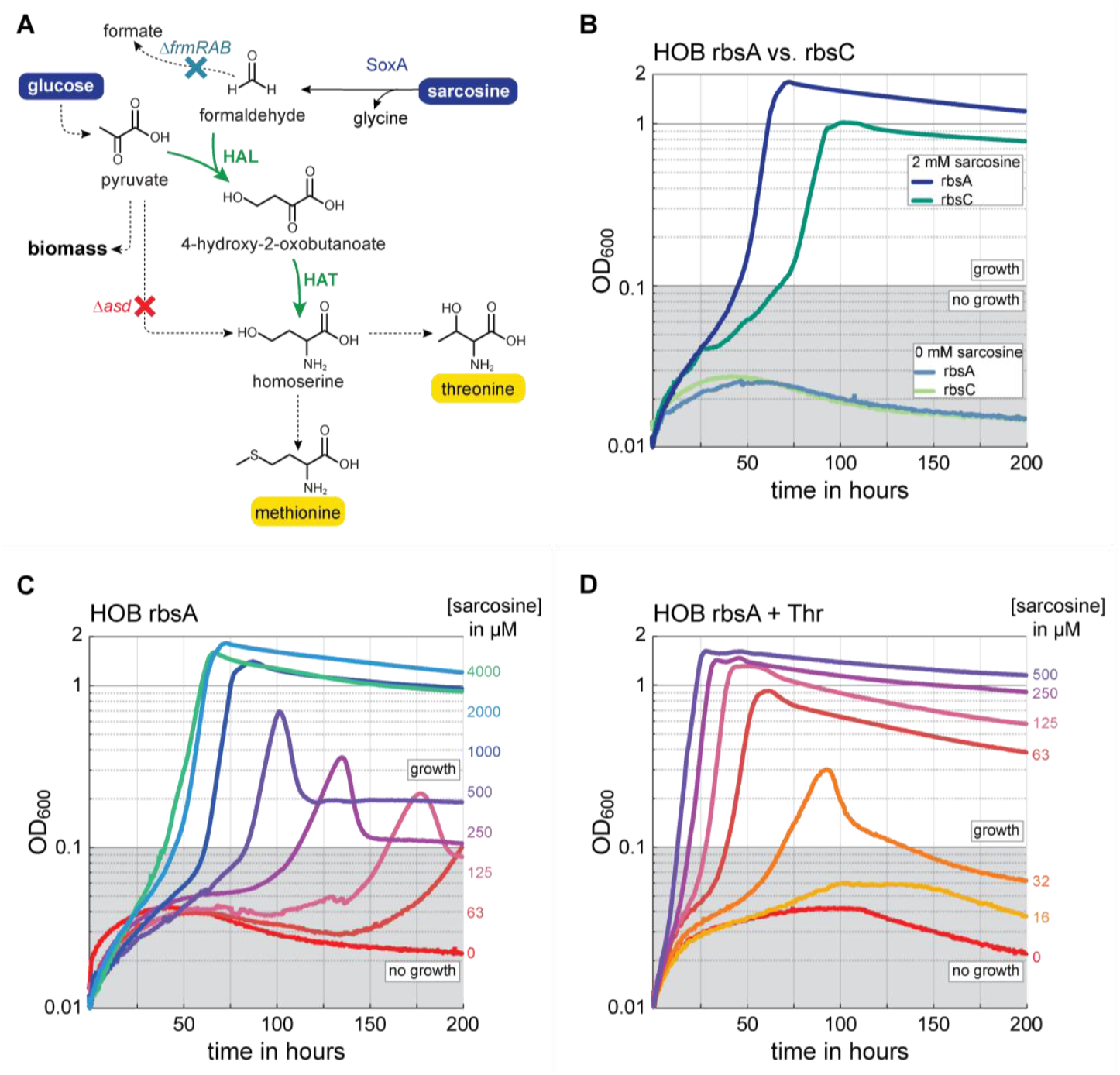
Addition of threonine enables the HOB biosensor to detect formaldehyde in the low micromolar range. (A) Metabolic scheme of the HOB biosensor. The strain is based on the promiscuous activity of HAL and HAT that condense formaldehyde and pyruvate into homoserine. In the HOB biosensor, the native homoserine biosynthesis pathway is disrupted by the deletion of aspartate semialdehyde dehydrogenase (*asd*), making the strain auxotrophic for threonine and methionine. These auxotrophies can be relieved upon the assimilation of intracellular formaldehyde by HAL and HAT. To increase the sensitivity of the strain, a formaldehyde sink was removed by deleting the endogenous formaldehyde detoxification system of *E. coli* (*frmRAB*). The HAL gene was expressed from the genome using a strong constitutive promoter and either a medium or strong ribosome binding site (rbsC or rbsA, respectively). SoxA was expressed from a plasmid to allow sarcosine conversion into formaldehyde. (B) The HOB biosensor with either rbsA or rbsC was cultivated in minimal medium, supplemented with 10 mM glucose, 1 mM isoleucine, 0.25 mM DAP with and without the addition of 2 mM sarcosine. (C) The HOB biosensor with rbsA was used for detailed characterization of the formaldehyde sensitivity. The strain was cultivated in a gradient of sarcosine concentrations ranging from 0 to 2000 μM. The experiment showed a correlation between growth and sarcosine concentration ranging from 125 to 2000 μM. (D) When 1 mM threonine was supplemented to reduce the formaldehyde dependency of the strain, the HOB biosensor was able to detect formaldehyde concentrations as low as 32 μM. Abbreviations: *asd*, aspartate semialdehyde dehydrogenase; *frmRAB*, formaldehyde detoxification system; SoxA, sarcosine oxidase; HAL, HOB aldolase; HAT, HOB amino transferase. Figure elements: yellow background: auxotrophy; blue circle: substrate; red cross: deletion causing auxotrophy; cyan cross: non-essential deletion; green arrow: formaldehyde assimilation flux; dashed arrow: multi enzymatic reaction.

To examine the strain’s suitability as a formaldehyde biosensor, we engineered the HOB biosensor by deleting *asd, frmRAB* and the endogenous *rhmA* gene. We then expressed *rhmA* with either a medium or a strong RBS (rbsC and rbsA, respectively) under the control of a constitutive promoter from a safe spot in the genome (Bassalo et al., 2016). The growth of both strains expressing SoxA from a plasmid was tested with a defined concentration of sarcosine (**Figure 4B**). As the expression of *rhmA* with rbsA resulted in better growth, this strain was used in all further experiments. Next, we performed a detailed characterization experiment with a sarcosine gradient to examine the sensitivity of the strain. The strain was able to grow with sarcosine concentrations as low as 125 μM making it more sensitive than the other two biosensors. As the formaldehyde sensitivity of the HOB biosensor could theoretically be further increased by supplementing either methionine or threonine (neither amino acid can be converted to homoserine, thus allowing supplementation without risking a complete relieve of the auxotrophies), we tested the addition of either amino acid to the selective medium. An initial screening experiment suggested that supplementation of methionine does not improve the sensitivity of the strain whereas supplementation of threonine does (**Figure S4**). Thus, we decided to investigate the formaldehyde sensitivity of the HOB biosensor with threonine supplementation. Notably, the addition of threonine reduces the formaldehyde dependency of the strain to 0.1539 mmol/gCDW (as estimated by FBA). By repeating the gradient experiment with supplementation of 1 mM threonine, the strain was capable of growing with sarcosine concentrations as low as 30 μM, making it by far the most sensitive biosensor suitable for the detection of low intracellular formaldehyde.

### Comparison of formaldehyde sensitivity reveals operational range of biosensor strains

After characterizing the biosensor strains in detail, we aimed to systematically compare their formaldehyde sensitivity and to determine their operational range. The operational range is the range of formaldehyde concentrations over which the biosensor exhibits a change in final OD_600_ (Rogers et al., 2016). To this end, we determined the maximum (max.) OD_600_ reached by the respective biosensor at a given sarcosine concentration and visualized the data in a sensitivity plot (**Figure 5A**). As for most of the strains the lag phase increased when lower sarcosine concentrations were used, growth starting after 200 hours was not considered. When comparing all strains, it becomes clear that the HOB biosensor with supplementation of threonine is the most sensitive strain followed by the LtaE^*^, LtaE and the RuMP sensor. As the operational ranges of the sensors overlap (especially for HOB, LtaE^*^ and LtaE), we suggest to use the formaldehyde biosensors in the following way: the HOB biosensor (with the addition of threonine) should be used for detecting formaldehyde concentrations between 30 - 200 μM, the LtaE^*^ biosensor for concentrations between 200 μM - 1 mM and the RuMP biosensor for concentrations between 1 - 10 mM (**Figure 5B**). In this combination, the formaldehyde biosensors present an operational range that spans almost three orders of magnitude, allowing for a wide range of applications.

**Figure 5:**
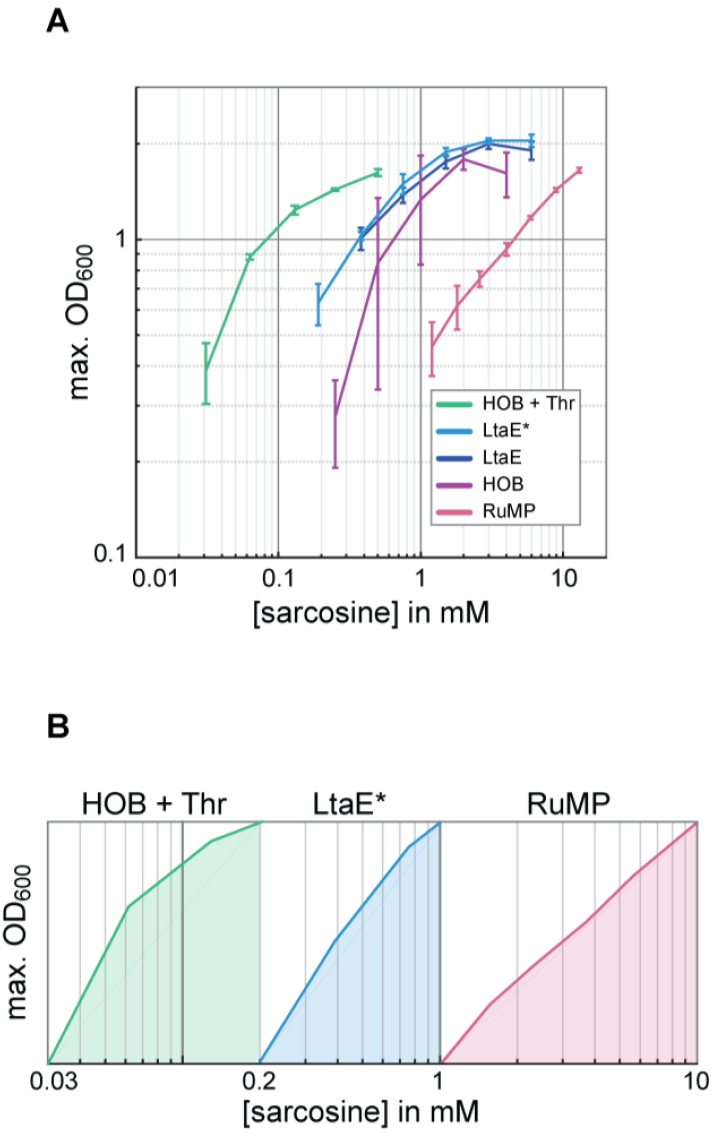
Comparison of formaldehyde sensitivities reveals the operational range of the biosensors. (A) To create a sensitivity plot that allows comparison of all formaldehyde biosensors, the maximum OD_600_ reached at specific a sarcosine concentration in the gradient experiments was plotted. While threonine supplementation in the HOB biosensor shows a strong increase on the final OD_600_, the evolved LtaE^*^ strain grows almost to the same max. OD_600_ but can grow with lower sarcosine concentrations. The RuMP biosensor requires the highest sarcosine concentration to reach its max. OD_600_ when compared to all other strains. (B) Together, the HOB biosensor (with addition of threonine), the LtaE^*^ biosensor and the RuMP biosensor present an operational range from 0.03 to 10 mM spanning almost 3 orders of magnitude. For each concentration of sarcosine, the average maximum OD_600_ reached of three independent growth experiments is shown and the standard deviations are reprented by the error bars.

### Direct addition of formaldehyde indicates potential of biosensors for environmental applications

The ability to detect extracellular formaldehyde would make the biosensors suitable for environmental or industrial applications. As such, they could be used to assess the formaldehyde level in a probe provided, if it does not contain any of the metabolites that would release the auxotrophy of the biosensor. To investigate the potential of the biosensors for such purposes, we tested growth of the strains upon direct addition of formaldehyde to the medium. To this end, a gradient of formaldehyde was tested. Surprisingly, both the HOB and the LtaE^*^ biosensors grew well upon direct addition of formaldehyde, showcasing almost the same sensitivity pattern as in the sarcosine experiments (**Figure 6**). Only the RuMP biosensor did not grow with direct addition of formaldehyde. We argue that this is related to the high *K*_*m*_ of HPS and the high amount of formaldehyde required to relieve the auxotrophies of the RuMP biosensor (>1 mM as approximated from the sarcosine gradient experiment). Thus, formaldehyde at concentrations that could relieve the auxotrophies becomes toxic preventing cell growth. This hypothesis is in accordance with literature data which suggests that formaldehyde becomes highly toxic to *E. coli* at concentrations >400 μM (Gutheil et al., 1997; Patterson et al., 2020). The results of these experiments unequivocally confirm the suitability of the different strains to serve as formaldehyde biosensors.

**Figure 6:**
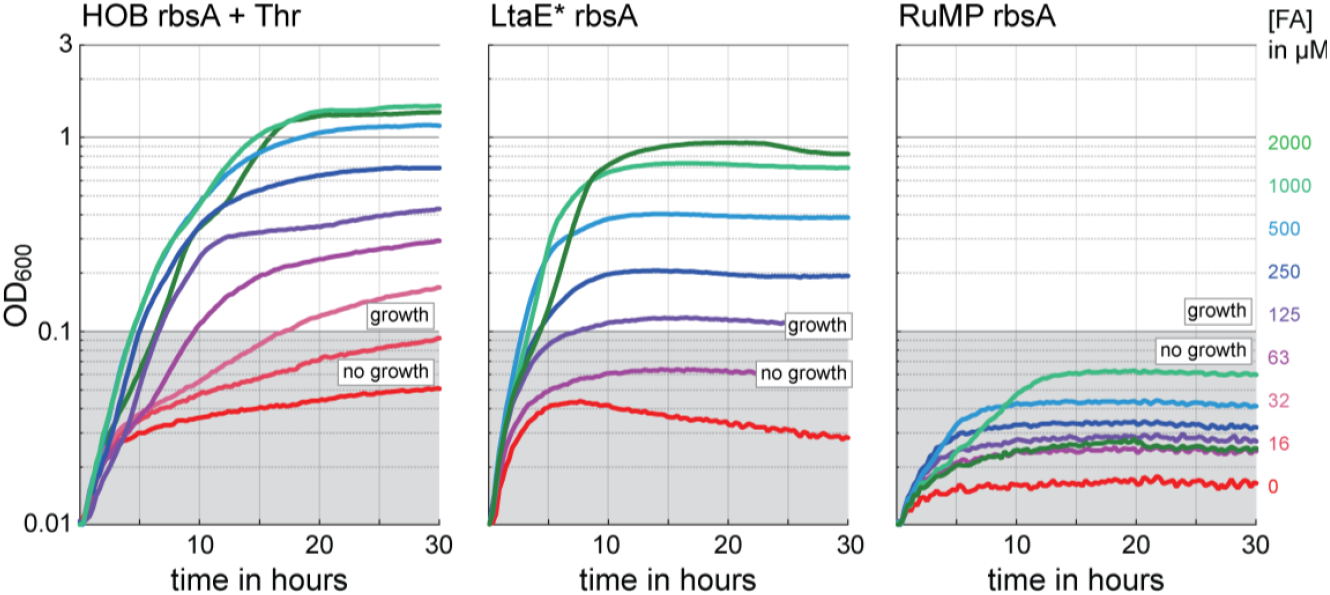
The LtaE^*^ and HOB biosensors can sense extracellular formaldehyde. To investigate the effect of directly added formaldehyde on the biosensors’ growth phenotype, the strains were tested in the respective medium (same as in sarcosine gradient except without sarcosine addition) with a formaldehyde gradient ranging from 0 to 2000 μM. The HOB biosensor reached its max. OD_600_ at a formaldehyde concentration of 1000 μM and could still grow with 32 μM. Also the LtaE^*^ biosensor grew upon direct supplementation of formaldehyde. Showing a concentration dependent growth phenotype, the strain was able to grow with formaldehyde concentrations ranging from 125 to 2000 μM. The RuMP biosensor was not able to grow upon direct addition of formaldehyde.

### Benchmarking of biotechnologically relevant MDHs in the HOB sensor

After determining the HOB biosensor with addition of threonine as the most sensitive formaldehyde sensor, we aimed to test its applicability for screening formaldehyde producing enzymes. This could be highly useful for enzyme engineering efforts or the discovery of novel formaldehyde producing enzymes. As in such cases rather low initial formaldehyde production rates are expected, we decided to screen the *in vivo* activity of widely used MDHs. To this end, we transformed the HOB biosensor with plasmids encoding *Bacillus stearothermophilus* MDH (*Bs*MDH), *C. glutamicum* MDH (*Cg*MDH), *Cupriavidus necator* N-1 MDH (*Cn*MDH) and an engineered *Bacillus methanolicus* MDH (*Bm*MDH^*^) and tested the strains growth with methanol as a formaldehyde source (**Figure 7A**). While all tested MDHs supported growth of the HOB biosensor at methanol concentrations of 80 mM and 40 mM, only the *Bs*MDH (which resulted in highest OD_600_ for all tested methanol concentrations) was able to provide sufficient formaldehyde to enable growth of the strain. Looking at the max. OD_600_ at different concentrations, it becomes clear that the *Bs*MDH is by far the best enzyme followed by the *Cn*MDH and *Cg*MDH. The experiment shows that the HOB biosensor is a suitable platform for testing the *in vivo* activity of formaldehyde generating enzymes.

**Figure 7:**
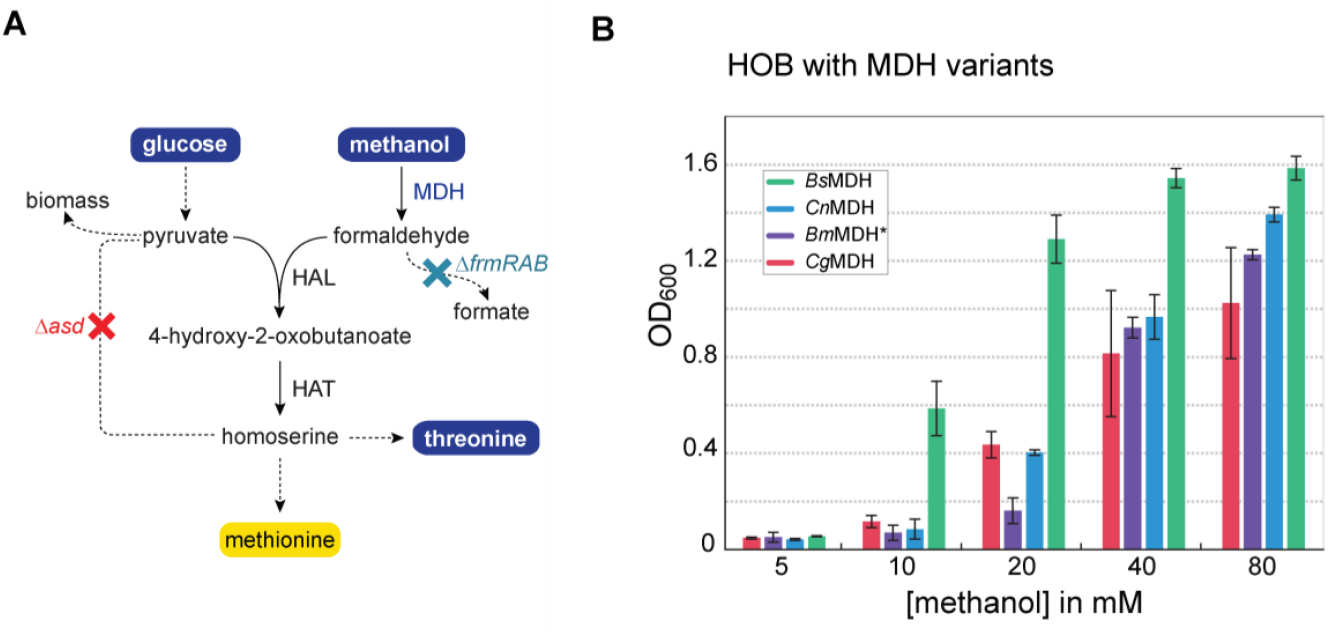
The HOB biosensor serves as an *in vivo* screening platform for formaldehyde producing enzymes. (A) MDH expression in the HOB biosensor allows methanol oxidation to formaldehyde and its subsequent assimilation into homoserine via the combined HAL and HAT activities. Different biotechnological relevant MDH variants were expressed in the HOB sensor from a plasmid using a strong constitutive promoter and a medium strength rbsC. (B) The HOB rbsA biosensor expressing either MDH variant was cultivated in minimal medium, supplemented with 10 mM glucose, 1 mM isoleucine, 1 mM threonine, 0.25 mM DAP and 5 to 80 mM methanol. While all MDH variants rescued growth at 80 mM methanol, only the *Bs*MDH was capable of rescuing growth at 10 mM methanol. Abbreviations: *asd*, aspartate semialdehyde dehydrogenase; *frmRAB*, formaldehyde detoxification system; *Bs*MDH, *Bacillus stearothermophilus* MDH; *Bm*MDH^*^, engineered *Bacillus methanolicus* MDH; *Cg*MDH, *Corynebacterium glutamicum* MDH; *Cn*MDH, *Cupriavidus necator* N-1 MDH; HAL, HOB aldolase; HAT, HOB amino transferase. Figure elements: yellow circle, auxotrophy; blue circle, substrate; red cross, deletion causing auxotrophy; cyan cross, non-essential deletion; dashed arrow, multi enzymatic reaction.

## Discussion

Cell-based biosensors have become an attractive, cost-effective tool in biotechnological applications ranging from environmental monitoring to clinical diagnosis (Gupta et al., 2019). Compared to chemical sensors, they are self-assembling, easy to handle, cheap and of low environmental impact. An ideal biosensor should be highly specific and detect a wide range of concentrations of the target molecule. Furthermore, it should have a high signal to noise ratio and a low false-positive rate (Dietrich et al., 2010). The concept of growth-coupled design has been recognized as a powerful methodology for the generation of cell-based biosensors that can detect key metabolites or enzyme activities with said characteristics (Aslan et al., 2020; Wenk et al., 2020).

In the present work, we employed a growth-coupled design strategy for the creation of three distinct formaldehyde biosensors that can detect a wide range of intracellularly produced formaldehyde concentrations spanning almost three orders of magnitude. By strategically interrupting central metabolic fluxes through gene deletions, we made the strains dependent on formaldehyde assimilation for cellular growth. Formaldehyde assimilation was enabled through genomic expression of formaldehyde assimilating enzymes and verified in various experiments using sarcosine as a thermodynamically and kinetically favorable source of formaldehyde. Sarcosine also known as N-methyl-glycine is a non-toxic amino acid, that can be efficiently cleaved to formaldehyde and glycine. It has been used in this and previous studies as a preferred formaldehyde source over the slow and limiting methanol oxidation or direct formaldehyde addition (He et al., 2020, 2018).

In a previous study, a highly sensitive plasmid-based formaldehyde biosensor that leads to GFP expression upon intracellular formaldehyde has been engineered (Woolston et al., 2018). This sensor showed an operational range of ∼1 - ∼250 μM formaldehyde. Thus, the growth biosensors presented in this work, which together present an operational range of ∼30 - ∼13 mM, ideally complement the previous work and dramatically increase the operational range for detecting formaldehyde by using *E. coli* biosensors. Compared to the GFP-based sensor, the growth biosensors constructed in the present work are fully genomic and produce an easy to detect signal: cell growth. Due to their stringent design, the sensor strains are metabolically robust allowing their use in long-term experiments such as ALE.

In the design phase of this study, we assessed the formaldehyde dependency of the biosensors via FBA in order to create three strains with different sensitivities. Our results suggested that the RuMP biosensor would be the most sensitive followed by the LtaE biosensor and the HOB biosensor. Since the calculations did not consider important cellular factors like co-factor availability or enzyme kinetics, it is of little surprise, that the experimental validation of the strains resulted in a different picture than the FBA prediction: While the engineered strains differ in their formaldehyde sensitivity, the characterization experiments suggest that the HOB biosensor is the most sensitive strain followed by LtaE and RuMP. This shows clearly that FBA is a good tool for *a priori* estimations but cannot replace experimental validations. As current models do not consider important factors like enzyme kinetics or co-factor availability, they can hardly predict the real sensitivity of a biosensor.

Due to the wide operational range, the three biosensors could be used for applications ranging from enzyme discovery to enzyme engineering. As showcased by the comparison of *in vivo* catalytic efficiencies of biotechnologically relevant MDHs, the sensor strains can be easily employed for screening libraries of formaldehyde producing enzymes. Here, a highly sensitive strain like the HOB biosensor could be used to identify activities in an unknown library (e.g. metagenomic library). Whereas, the less sensitive strains could be used for comparing enzyme efficiencies of engineered enzymes that are known to already have a basal activity. Ultimately, using the RuMP biosensor, one could even select for the generation of >70 % biomass through the enzyme of interest. The design of the strain allows cultivation on xylose and a formaldehyde source alone (leaving out succinate as a substrate for lower metabolism), which selects for the generation off all biomass except the pentose sugars from Ru5P and formaldehyde.

Another potential area of application could be the testing of environmental or industrial probes for their formaldehyde content. As both the LtaE^*^ and the HOB biosensor were able to grow upon direct addition of formaldehyde to the medium, the biosensors could be used to detect formaldehyde contamination or provide an estimate of the formaldehyde concentration of a solution. It should be considered, that the range of direct formaldehyde detection is limited (125 - 2000 μM) and that contaminations with amino acids (specifically serine, homoserine, threonine and methionine) could lead to false positives as they would directly release the auxotrophies of the strains.

Taken together, the formaldehyde biosensors created in this work, present an ideal tool for enzyme discovery and enzyme engineering studies that aim to find or improve formaldehyde producing enzymes. The strains might also be used for environmental or industrial applications; however, further investigations of their applicability in these areas are still required.

## Supporting information

Gene sequences

## Acknowledgements

This work was supported by the German Ministry of Education and Research Grant 031B0850B (MetAFor) and the Max Planck Society. The authors are grateful to Arren Bar-Even who initiated the project and was an inspiring mentor to all of us. The authors thank Nicole Paczia and Peter Claus for performing the HPLC-MS measurements and Enrico Orsi for critical reading of the manuscript and helpful suggestions.

## Author contributions

S.W. and A.B.-E. conceptualized the project and designed and supervised the research. S.W. and K.S. wrote the manuscript with contributions from J.B., M.B., M.N. and H.H. S.W., K.S., J.B. and M.B. designed and performed the experiments and analyzed the data. K.S., J.B., M.B, P.K. and H.H. genetically engineered the *E. coli* biosensor strains. T.J.E. supervised the *in vitro* work. M.N. performed the enzymatic assays for this study. H.H. performed flux balance analysis.

## Methods

### Chemicals and reagents

Primers were synthesized at Integrated DNA Technologies (IDT). DNA fragments were amplified with the high-fidelity polymerase PrimeSTAR Max DNA Polymerase from TaKaRa and colony PCR reactions were performed using DreamTaq polymerase (Thermo Fisher Scientific). FastDigest restriction enzymes and T4 DNA ligase from Thermo Fisher Scientific were used for restriction-based cloning. The kits used for plasmid isolation and PCR cleanup were purchased from by Thermo Fischer Scientific. All chemicals used in the plate reader experiments were ordered from Sigma-Aldrich.

### Growth media

For cloning and strain engineering, cultures were grown in lysogeny broth (LB) medium (10 g/L NaCl, 10 g/L Tryptone, 5 g/L Yeast Extract) supplemented with the relevant antibiotic (50 μg/mL kanamycin, 30 μg/mL chloramphenicol or 100 μg/mL streptomycin). Diaminopimelate (DAP, 0.25 mM) needs to be supplemented to the growth media of the HOB strain due to the asd knockout. During growth experiments, cells were cultivated in M9 minimal medium (50 mM Na2HPO4, 20 mM KH2PO4, 1 mM NaCl, 20 mM NH4Cl, 2 mM MgSO4 and 100 μM CaCl2), supplemented with trace elements (134 μM EDTA, 31 μM FeCl3, 6.2 μM ZnCl2, 0.76 μM CuCl2, 0.42 μM CoCl2, 1.62 μM H3BO3, 0.081 μM MnCl2) and the relevant carbon sources.

### Strains

Plasmid construction was done in *E. coli* DH5α. The *E. coli* SIJ488 strain, which is derived from K-12 MG1655, was used as a base strain for the biosensor strains (Jensen et al., 2015). The KEIO collection was used as donor strains to prepare the P1 donor lysates for phage transduction (Baba et al., 2006). The strains used in this study are listed in Supplementary Table 1.

### Strain engineering

The biosensor strains were constructed by using recombineering techniques to modify the SIJ488 base strain. Deletions were made by P1 phage transduction and λ-Red recombineering as described in (Wenk et al., 2018). To make deletions with λ-Red recombineering, a kanamycin or chloramphenicol antibiotic cassette, flanked by FRT sites, was amplified with primers containing 50bp homology arms that are flanking the region of the target gene. This linear DNA fragment was introduced in the cell by electroporation after one hour of inducing the λ-Red recombineering machinery with 15 mM arabinose. The cell suspension was then plated on LB plates containing the relevant antibiotics and a colony PCR was performed to confirm successful gene deletions. The antibiotics cassettes were removed by inducing the flippase with 50 μM L-rhamnose for at least four hours and removal was confirmed by a colony PCR. For P1 phage transduction, the KEIO collection strain with the desired knockout was grown in LB supplemented with 5 mM CaCl_2_ until an OD ∼0.3. Then the culture was infected with P1vir (Ikeda and Tomizawa, 1965) stock lysate for 2-4 hours. The cells were pelleted by centrifugation and the supernatant was filtered through a 0.22-μm filter to obtain the donor lysate that was then used for the transduction. The deletion in the relevant strain was made by resuspending the pellet of an overnight culture in P1 salts. P1 donor lysate was added to this cell suspension and incubated for 30 min at 30°C. After a recovery for 1 hour in LB with ∼100 mM sodium citrate, the cells were plates on selective plates containing 5 mM sodium citrate and kanamycin. The antibiotics cassette was later removed by inducing the flippase. Knockouts and removal of the antibiotics cassette were confirmed by a colony PCR.

Genomic integrations were made via a conjugation-based genetic recombination method as described in (Wenk et al., 2018). In short, the synthetic operon was cloned into a vector which contains the chloramphenicol resistance gene (cam^R^) and 600bp homology arms that are compatible with the genomic integration site. Furthermore, the vector contains a levansucrase gene (sacB) and the traJI gene to allow for the transfer of the plasmid. After transformation in chemically competent ST18, the colonies growing on chloramphenicol plates were screened by a colony PCR. A positive clone was then used for the conjugation. After conjugation, the SIJ488 strains growing on chloramphenicol plates were screened by sucrose counter selection and kanamycin resistance tests to isolate recombinant SIJ488 strains with the correctly integrated synthetic operon. The antibiotics cassette was removed by inducing the flippase. The genomic integration and antibiotics cassette removal were confirmed by a colony PCR.

### Synthetic-Operon construction

All genes were codon optimized for *E. coli* K-12, after which an 6xHis-tag was added to the N-terminal side. Subsequently, the genes were inserted into a cloning vector, which attaches a synthetic ribosomal binding site upstream of the gene (Zelcbuch et al., 2013). These cloning vectors were used to assemble synthetic operons via restriction and ligation using the protocol as described in (Wenk et al., 2018).

The assembled construct was then transferred to the pZ expression vector which contains the p15A ori, a streptomycin resistance gene and attaches the Ppgi-20 promoter to the gene/operon of interest. This assembly was performed using the restriction enzymes EcoRI and PstI. (Wenk et al., 2018).

### Growth experiments

Prior to all growth experiments, the strains were streaked out from the glycerol stock on LB plates (supplemented with the respective antibiotic and 0.25 mM DAP in case of the HOB biosensor) and grown overnight at 37°C. From these plates, precultures in relaxing liquid medium (as indicated in Table 1) were inoculated for overnight growth at 37°C and 220 rpm. The cells were harvested (11000 g, 1 min), washed three times with M9 mineral medium, and diluted to a final OD_600_ of 0.01 in the desired medium. In a Nunc 96-well microplates (Thermo Fisher Scientific), 150 μl of the culture was added to each well and the wells were covered with 50 μl of mineral oil (Sigma-Aldrich) to prevent evaporation. The growth experiments were performed in the BioTek Epoch2 plate reader (BioTek Instrument, USA) which measured the optical density after each kinetic cycle. A cycle consists of 12 shaking steps in which 60 seconds linear and 60 seconds orbital shaking were alternated (1 mm amplitude). The OD_600_ was corrected to present the OD_600_cuvette_ by dividing the OD_600_plate_ by the correction factor 0.23. The mean of the technical duplicates is shown in the growth curves. The sensitivity plots show the average values of three independent experiments and their respective standard deviation. Data was only included when all three experiments showed growth.

**Table 1:**
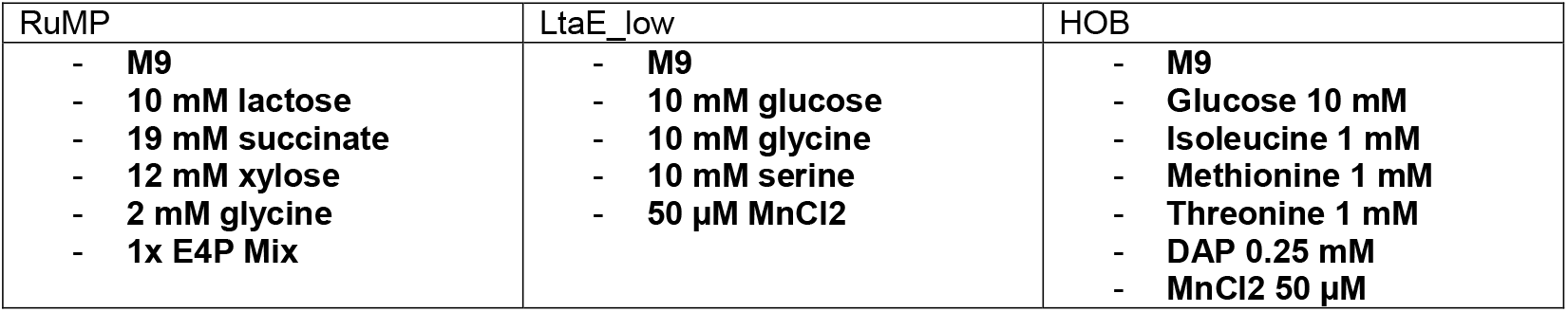
Composition of relaxing media for the different formaldehyde biosensors

| RuMP | LtaE_low | HOB |
| --- | --- | --- |
| - **M9**<br>- **10 mM lactose**<br>- **19 mM succinate**<br>- **12 mM xylose**<br>- **2 mM glycine**<br>- **1x E4P Mix** | - **M9**<br>- **10 mM glucose**<br>- **10 mM glycine**<br>- **10 mM serine**<br>- **50 µM MnCl<sub>2</sub>** | - **M9**<br>- **Glucose 10 mM**<br>- **Isoleucine 1 mM**<br>- **Methionine 1 mM**<br>- **Threonine 1 mM**<br>- **DAP 0.25 mM**<br>- **MnCl<sub>2</sub> 50 µM** |

#### Adaptive laboratory evolution of the LtaE biosensor

The experiment was conducted with the rbsA LtaE biosensor strain expressing CgMDH from a plasmid. Four 14 ml glass tubes with biological replicates were cultivated in selective medium (M9 minimal medium with 10 mM glucose, 10 mM glycine, 50 μM MnCl2 and 500 – 10 mM methanol) for 15 weeks during which the methanol concentration was stepwise reduced. Whenever the strain reached an OD_600_ > 0.5 the culture was diluted into fresh medium to an OD_600_ of 0.01. First, the strains were cultivated for one generation with 500 mM methanol. Then the methanol concentration was reduced to 100 mM for another generation. Thereafter the culture was diluted into medium containing 50 mM methanol. After 10 days, the methanol concentration was reduced to 25 mM. In the following, the methanol concentration was reduced to 20 mM, 15 mM and finally to 10 mM changing the methanol concentration approximately every 10 days. Growth in selective medium with 10 mM methanol was continued for 4 weeks. Then, single colonies were isolated from the growing populations and analysed via growth experiments and whole genome sequencing.

#### Whole-genome sequencing

Genomic DNA was extracted using a commercial kit (Mackerey Nagel) starting from 2-4 ml of overnight culture in LB. A minimum of 300 ng was sent for sequencing to NovoGene. Results were analyzed using the open source breseq software (Barrick et al., 2014). All NGS data was deposited at the Genome Sequence Archive.

### In vitro characterization of LtaE and LtaE^*^

Kinetics for LtaE and LtaE^*^ were determined by HPLC-MS detection of serine. Reaction mixtures contained 100 mM KxHyPO4, 5 mM MgCl_2_, 100 μM pyridoxal phosphate, 1 μM enzyme and the glycine and formaldehyde concentrations provided in Table SIX. After incubation at 30 °C (timepoints 0.5, 1, 2, and 3 min), samples were quenched in formic acid to a final concentration of 5 %. A serine standard in reaction matrix was run alongside the samples. Serine was detected via HPLC-MS and quantified against the standard. Initial slope of serine production was calculated by linear fit in GraphPad Prism v9 and Michaelis Menten kinetics were constructed in the same program using standard setting.

### HPLC-MS detection of amino acids

Quantitative determination of serine and glycine was performed using a LC-MS/MS. The chromatographic separation was performed on an Agilent Infinity II 1290 HPLC system using a ZicHILIC SeQuant column (150 × 2.1 mm, 3.5 μm particle size, 100 Å pore size) connected to a ZicHILIC guard column (20 × 2.1 mm, 5 μm particle size) (Merck KgAA) a constant flow rate of 0.3 ml/min with mobile phase A being 0.1 % Formic acid in 99:1 water:acetonitrile (Honeywell, Morristown, New Jersey, USA) and phase B being 0.1 % formic acid 99:1 acetonitrile:water (Honeywell, Morristown, New Jersey, USA) at 25° C.

The injection volume was 1 μl. The mobile phase profile consisted of the following steps and linear gradients: 0 – 5 min from 80 to 65 % B; 5 – 7 min from 65 to 20 % B; 7 – 9 min constant at 20 % B; 9 – 10 min from 20 to 80 % B; 10 to 12 min constant at 80 % B. An Agilent 6495 ion funnel mass spectrometer was used in positive mode with an electrospray ionization source and the following conditions: ESI spray voltage 2000 V, nozzle voltage 1000 V, sheath gas 250° C at 12 l/min, nebulizer pressure 60 psig and drying gas 100° C at 11 l/min. Compounds were identified based on their mass transition and retention time compared to standards. Chromatograms were integrated using MassHunter software (Agilent, Santa Clara, CA, USA). Absolute concentrations were calculated based on an external calibration curve prepared in the sample matrix.

Mass transitions, collision energies, Cell accelerator voltages, and Dwell times have been optimized using chemically pure standards. The parameter settings of all targets are given in the table below.

**Table 2:** Parameter settings of HPLC-MS targets

| Compound | Precursor | Product | Dwell time | Fragmenter Voltage | Collision Energy | Cell Accelerator Voltage | Polarity |
| --- | --- | --- | --- | --- | --- | --- | --- |
| Serin | 106.1 | 60.2 | 200 | 380 | 12 | 5 | Pos |
| Serin | 106.1 | 42.1 | 200 | 380 | 24 | 5 | Pos |
| Glycine | 76 | 76 | 200 | 380 | 0 | 5 | Pos |
| Glycine | 76 | 30.1 | 200 | 380 | 5 | 5 | Pos |

### Flux balance analysis

Flux balance analysis (FBA) was used for calculating formaldehyde dependency of the biosensor strains. The modeling was conducted with COBRApy (Ebrahim et al., 2013) using the most updated *E. coli* genome-scale metabolic model *i*ML1515 (Monk, 2017) with curations and changes. The lowest possible fluxes of formaldehyde uptake in the biosensor strains for different growth rates were calculated. Formaldehyde dependency of the biosensors was estimated from the slope of growth rates and minimal formaldehyde uptake rates from the modeling. The full code, including the construction of models for selection strains, was deposited at Zenodo.

## Data availability

Additional information on the experimental setup as well as detailed results are available from the corresponding author upon request. Any strains and plasmids generated during this study are available upon completing a Materials Transfer Agreement.

## Code availability

MATLAB and breseq codes used for the analysis of the experiments are available from the corresponding author upon request. The flux balance analysis code is available openly on Zenodo with DOI of 10.5281/zenodo.8009619.

