## Supplementary material for "Design, construction and optimization of formaldehyde growth biosensors with broad application in Biotechnology": Table S

### Supplementary Table 1: Strains used in this study

| Strain | Genotype | Source |
|---|---|---|
| DH5α | $F^- \lambda^-$ *Φ80lacZΔM15 Δ(lacZYA-argF)U169 deoR recA1 endA1 hsdR17*($rK^-$ $mK^+$) *phoA supE44 thi-1 gyrA96 relA1* | (Hanahan, 1983) |
| ST18 | *pro thi* $hsdR^+$ $Tp^r$ $Sm^r$; *chromosome::RP4-2 Tc::Mu-Kan::Tn7/λpir ΔhemA* | (Thoma and Schobert, 2009) |
| MG1655 | $F^- \lambda^-$ $ilvG^-$ *rfb-50 rph-1* | (Blattner et al., 1997) |
| SIJ488 | MG1655 *Tn7::para-exo-beta-gam; prha-FLP; xylSpm-IsceI* | (Jensen et al., 2015) |
| RuMP parent strain | | This study |
| RuMP biosensor (rbsA) | | This study |
| RuMP biosensor (rbsC) | | This study |
| LtaE parent strain | SIJ488 ΔfrmRAB ΔserA ΔglyA | This study |
| LtaE biosensor (rbsA) | SIJ488 ΔfrmRAB ΔpatZ ΔserA ΔglyA ΔltaE SS9:Ps:$RBS_A$:ltaE | This study |
| LtaE biosensor (rbsB) | SIJ488 ΔfrmRAB ΔpatZ ΔserA ΔglyA ΔltaE SS9:Ps:$RBS_C$:ltaE | This study |
| HOB parent strain | SIJ488 *ΔfrmRAB Δasd* | (He et al., 2020) |
| HOB biosensor (rbsA) | SIJ488 ΔackA-pta ΔfrmRAB ΔpatZ Δasd ΔrhmA SS2:Ps:$RBS_A$:rhmA | This study |
| HOB biosensor (rbsC) | SIJ488 ΔackA-pta ΔfrmRAB ΔpatZ Δasd ΔrhmA SS2:Ps:$RBS_C$:rhmA | This study |

### Supplementary Table 2: Plasmids used in this study

| Plasmid | Genes | Source | Uniprot/GenBank |
|---|---|---|---|
| pZ:ASS | p15A ori; $Strep^R$; $P_{pgi-20}$ | Lab collection | |
| pZ:ASS-SoxA | pZ:$P_{strong}$:$RBC_C$:SoxA | (He et al., 2018) | P40859 |
| pZ:ASS:C:cgMDH | pZ:$P_{strong}$:$RBC_C$:CgMDH | (Wenk et al., 2020) | A4QHJ5 |
| pZ:ASS:C:BsMDH | pZ:$P_{strong}$:$RBC_C$:BsMDH | (Wenk et al., 2020) | P42327 |
| pZ:ASS:C:BmMDH* | pZ:$P_{strong}$:$RBC_C$:BmMDH* | (Wenk et al., 2020) | I3DVX6 |
| pZ:ASS:C:CnMDH | pZ:$P_{strong}$:$RBC_C$:CnMDH | (Wenk et al., 2020) | F8GNE5 |
