## Supplementary material for "Design, construction and optimization of formaldehyde growth biosensors with broad application in Biotechnology": Figure S

### Supplementary Figures

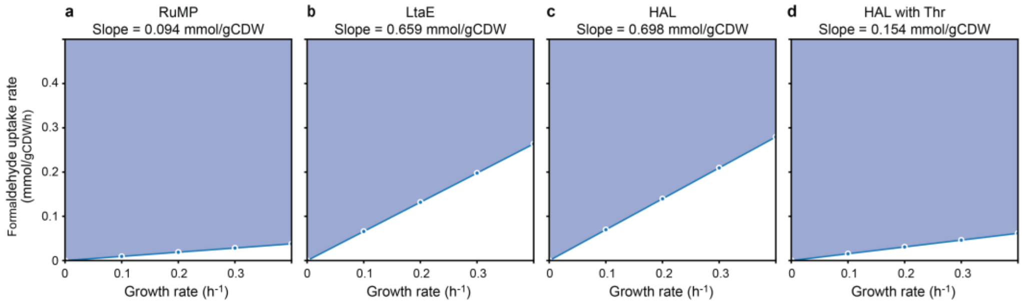


**Supplementary Figure S1: FBA estimations of formaldehyde dependency for different selection strains.**
FBA analysis was conducted using *E. coli* genome-scale model with relevant modifications. Minimal required formaldehyde uptake rates were calculated at different growth rates. The formaldehyde dependency of the strains was deduced from the slopes between growth rates and formaldehyde uptake rates. See Methods section for details.

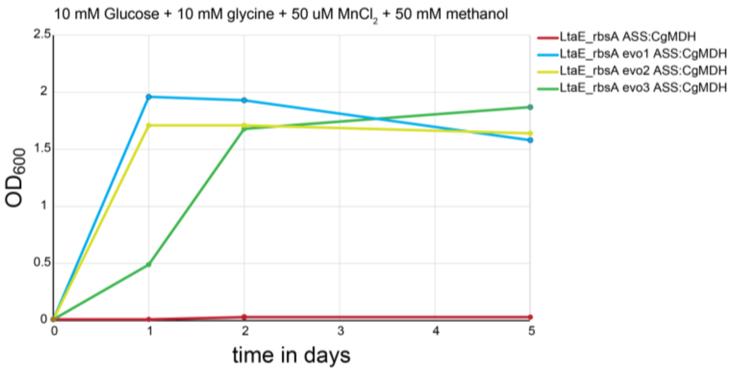


**Supplementary Figure S2: Comparison of evolved and unevolved LtaE biosensors**
Three single colonies isolated from the short tube ALE experiment (LtaE_rbsA evo1, evo2 and evo3) and the parent strain (LtaE_rbsA) were cultivated in closed 14 mL tubes in minimal medium supplemented with 50 mM methanol. Only evolved strains were able to grow. Genomic analysis of the evolved strains revealed a dedicated point mutation in the ltaE gene of all evolved strains which increases the catalytic efficiency of the enzyme.

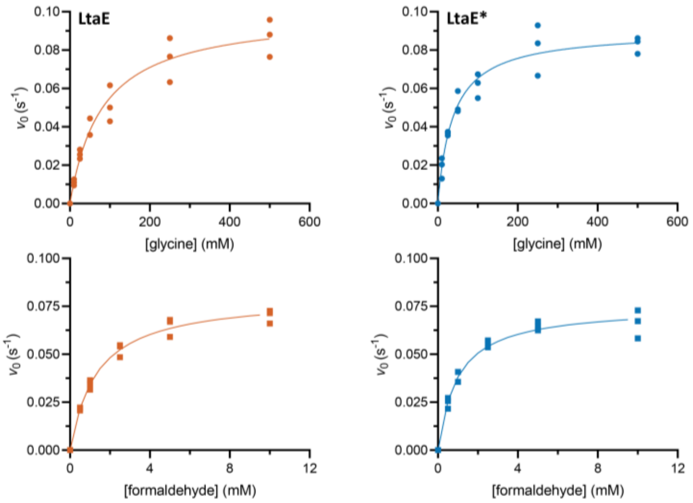


| glycine kinetics | |
|---|---|
| [glycine] (mM) | [formaldehyde] (mM) |
| 500, 250, 100, 50, 25, 10 | 5 |

| formaldehyde kinetics | |
|---|---|
| [glycine] (mM) | [formaldehyde] (mM) |
| 200 | 10, 5, 2.5, 1, 0.5 |

| enzyme | glycine | | | formaldehyde | | |
|---|---|---|---|---|---|---|
| | $k_{cat}$ ($s^{-1}$) | $K_m$ (mM) | $k_{cat}/K_m$ ($M^{-1}s^{-1}$) | $k_{cat}$ ($s^{-1}$) | $K_m$ (mM) | $k_{cat}/K_m$ ($M^{-1}s^{-1}$) |
| LtaE | 0.10 ± 0.00 | 81 ± 13 | 1.2 | 0.09 ± 0.00 | 1.3 ± 0.1 | 69 |
| LtaE* (C188Y) | 0.09 ± 0.00 | 38 ± 5 | 2.4 | 0.07 ± 0.00 | 1.0 ± 0.1 | 70 |

**Supplementary Figure S3: Kinetic evaluation of LtaE variants.**
Depicted are Michaelis Menten fits of initial velocities determined by HPLC-MS. Individual, independent replicates are shown (n = 3). Details of the experimental setup are given in the method section.

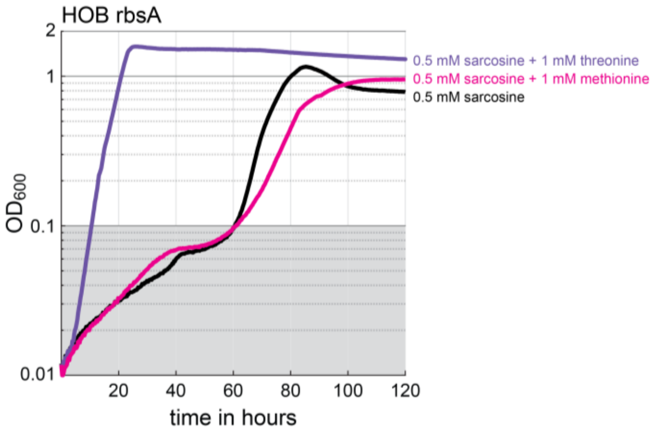


**Supplementary Figure S4: Addition of threonine improves sensitivity of the HOB biosensor**
The HOB biosensor expressing HAL with a strong rbsA and a sarcosine oxidase from a plasmid was cultivated in selective medium supplemented with 0.5 mM sarcosine +/- 1 mM of the amino acids threonine or metionine. As shown in the figure, only the addition of threonine improved the growth of the strain.
