## Supplementary material for "Design, construction and optimization of formaldehyde growth biosensors with broad application in Biotechnology": Gene sequences

**Codon optimized gene sequences (5’ -> 3’). 6×His tags are underlined.**

### 3-hexulose-6-phosphate synthase (HPS) from *Bacillus methanolicus* MGA3 (UniProt:I3DZR0)

ATGCATCATCACCATCACCACGAACTGCAACTGGCTCTGGACCTGGTTAACATCGAAGAAGCTAAACAGGTTGTTGCTGAAGTT
CAGGAATACGTTGACATCGTTGAAATCGGTACCCCGGTTATCAAAATCTGGGGTCTGCAAGCTGTTAAAGCTGTTAAAGACGCT
TTCCCGCATCTGCAAGTTCTGGCTGACATGAAAACCATGGACGCTGCTGCTTACGAAGTTGCTAAAGCTGCTGAACACGGTGC
TGACATCGTTACCATCCTGGCTGCTGCTGAAGACGTTTCTATCAAAGGTGCTGTTGAAGAAGCTAAAAAACTGGGTAAAAAAAT
CCTGGTTGACATGATCGCTGTTAAAAACCTGGAAGAACGTGCTAAACAGGTTGACGAAATGGGTGTTGACTACATCTGCGTTCA
CGCTGGTTACGACCTGCAAGCTGTTGGTAAAAACCCGCTGGACGACCTGAAACGTATCAAAGCTGTTGTTAAAAACGCTAAAA
CCGCTATCGCTGGTGGTATCAAACTGGAAACCCTGCCGGAAGTTATCAAAGCTGAACCGGACCTGGTTATCGTTGGTGGTGGT
ATCGCTAACCAGACCGACAAAAAAGCTGCTGCTGAAAAAATCAACAAACTGGTTAAACAGGGTCTGTAA

### 6-phospho-3-hexuloisomerase (PHI) from *Bacillus methanolicus* MGA3 (UniProt:Q6TV53)

ATGCATCATCACCATCACCACATCTCTATGCTGACCACCGAATTTCTGGCTGAAATCGTTAAAGAACTGAACTCTTCTGTTAACC
AGATCGCTGACGAAGAAGCTGAAGCTCTGGTTAACGGTATCCTGCAATCTAAAAAAGTTTTCGTTGCTGGTGCTGGTCGTTCTG
GTTTCATGGCTAAATCTTTCGCTATGCGTATGATGCACATGGGTATCGACGCTTACGTTGTTGGTGAAACCGTTACCCCGAACT
ACGAAAAAGAAGACATCCTGATCATCGGTTCTGGTTCTGGTGAAACCAAATCTCTGGTTTCTATGGCTCAGAAAGCTAAATCTA
TCGGTGGTACCATCGCTGCTGTTACCATCAACCCGGAATCTACCATCGGTCAGCTGGCTGACATCGTTATCAAAATGCCGGGT
TCTCCGAAAGACAAATCTGAAGCTCGTGAAACCATCCAGCCGATGGGTTCTCTGTTCGAACAGACCCTGCTGCTGTTCTACGA
CGCTGTTATCCTGCGTTTCATGGAAAAAAAAGGTCTGGACACCAAAACCATGTACGGTCGTCACGCTAACCTGGAATAA

### Sarcosine oxidase from *Bacillus sp. B-0618* (UniProt: P40859)

ATGCATCATCACCATCACCACTCTACCCACTTCGACGTTATCGTTGTTGGTGCTGGTTCTATGGGTATGGCTGCTGGTTACCAG
CTGGCTAAACAGGGTGTTAAAACCCTGCTGGTTGACGCTTTCGACCCGCCGCACACCAACGGTTCTCACCACGGTGACACCC
GTATCATCCGTCACGCTTACGGTGAAGGTCGTGAATACGTTCCGCTGGCTCTGCGTTCTCAGGAACTGTGGTACGAACTGGAA
AAAGAAACCCACCACAAAATCTTCACCAAAACCGGTGTTCTGGTTTTCGGTCCGAAAGGTGAATCTGCTTTCGTTGCTGAAACC
ATGGAAGCTGCTAAAGAACACTCTCTGACCGTTGACCTGCTGGAAGGTGACGAAATCAACAAACGTTGGCCGGGTATCACCGT
TCCGGAAAACTACAACGCTATCTTCGAACCGAACTCTGGTGTTCTGTTCTCTGAAAACTGCATCCGTGCTTACCGTGAACTGGC
TGAAGCTCGTGGTGCTAAAGTTCTGACCCACACCCGTGTTGAAGACTTCGACATCTCTCCGGACTCTGTTAAAATCGAAACCG
CTAACGGTTCTTACACCGCTGACAAACTGATCGTTTCTATGGGTGCTTGGAACTCTAAACTGCTGTCTAAACTGAACCTGGACA
TCCCGCTGCAACCGTACCGTCAGGTTGTTGGTTTCTTCGAATCTGACGAATCTAAATACTCTAACGACATCGACTTCCCGGGTT
TCATGGTTGAAGTTCCGAACGGTATCTACTACGGTTTCCCGTCTTTCGGTGGTTGCGGTCTGAAACTGGGTTACCACACCTTCG
GTCAGAAAATCGACCCGGACACCATCAACCGTGAATTTGGTGTTTACCCGGAAGACGAATCTAACCTGCGTGCTTTCCTGGAA
GAATACATGCCGGGTGCTAACGGTGAACTGAAACGTGGTGCTGTTTGCATGTACACCAAAACCCTGGACGAACACTTCATCAT
CGACCTGCACCCGGAACACTCTAACGTTGTTATCGCTGCTGGTTTCTCTGGTCACGGTTTCAAATTCTCTTCTGGTGTTGGTGA
AGTTCTGTCTCAGCTGGCTCTGACCGGTAAAACCGAACACGACATCTCTATCTTCTCTATCAACCGTCCGGCTCTGAAAGAATC
TCTGCAAAAAACCACCATCTAA

Methanol dehydrogenase from *Bacillus stearothermophilus* (UniProt: P42327)

ATGCATCATCACCATCACCACAAAGCTGCTGTTGTTAACGAATTTAAAAAAGCTCTGGAAATCAAAGAAGTTGAACGTCCGAAA
CTGGAAGAAGGTGAAGTTCTGGTTAAAATCGAAGCTTGCGGTGTTTGCCACACCGACCTGCACGCTGCTCACGGTGACTGGC
CGATCAAACCGAAACTGCCGCTGATCCCGGGTCACGAAGGTGTTGGTATCGTTGTTGAAGTTGCTAAAGGTGTTAAATCTATCA
AAGTTGGTGACCGTGTTGGTATCCCGTGGCTGTACTCTGCTTGCGGTGAATGCGAATACTGCCTGACCGGTCAGGAAACCCTG
TGCCCGCACCAGCTGAACGGTGGTTACTCTGTTGACGGTGGTTACGCTGAATACTGCAAAGCTCCGGCTGACTACGTTGCTAA
AATCCCGGACAACCTGGACCCGGTTGAAGTTGCTCCGATCCTGTGCGCTGGTGTTACCACCTACAAAGCTCTGAAAGTTTCTG
GTGCTCGTCCGGGTGAATGGGTTGCTATCTACGGTATCGGTGGTCTGGGTCACATCGCTCTGCAATACGCTAAAGCTATGGGT
CTGAACGTTGTTGCTGTTGACATCTCTGACGAAAAATCTAAACTGGCTAAAGACCTGGGTGCTGACATCGCTATCAACGGTCTG
AAAGAAGACCCGGTTAAAGCTATCCACGACCAGGTTGGTGGTGTTCACGCTGCTATCTCTGTTGCTGTTAACAAAAAAGCTTTC
GAACAGGCTTACCAGTCTGTTAAACGTGGTGGTACCCTGGTTGTTGTTGGTCTGCCGAACGCTGACCTGCCGATCCCGATCTT
CGACACCGTTCTGAACGGTGTTTCTGTTAAAGGTTCTATCGTTGGTACCCGTAAAGACATGCAGGAAGCTCTGGACTTCGCTG
CTCGTGGTAAAGTTCGTCCGATAGTTGAAACCGCTGAACTGGAAGAAATCAACGAAGTTTTCGAACGTATGGAAAAAGGTAAAA
TCAACGGTCGTATCGTTCTGAAACTGAAAGAAGACTAA

Methanol dehydrogenase from *Corynebacterium glutamicum* (Unitprot: A4QHJ5)

ATGCATCATCACCATCACCACACCACCGCTGCTCCGCAGGAATTTACCGCTGCTGTTGTTGAAAAATTCGGTCACGAAGTTACC
GTTAAAGACATCGACCTGCCGAAACCGGGTCCGAACCAGGCTCTGGTTAAAGTTCTGACCTCTGGTATCTGCCACACCGACCT
GCACGCTCTGGAAGGTGACTGGCCGGTTAAACCGGAACCGCCGTTCGTTCCGGGTCACGAAGGTGTTGGTGAAGTTGTTGAA
CTGGGTCCGGGTGAACACGACGTTAAAGTTGGTGACATCGTTGGTAACGCTTGGCTGTGGTCTGCTTGCGGTACCTGCGAATA
CTGCATCACCGGTCGTGAAACCCAGTGCAACGAAGCTGAATACGGTGGTTACACCCAGAACGGTTCTTTCGGTCAGTACATGC
TGGTTGACACCCGTTACGCTGCTCGTATCCCGGACGGTGTTGACTACCTGGAAGCTGCTCCGATCCTGTGCGCTGGTGTTACC
GTTTACAAAGCTCTGAAAGTTTCTGAAACCCGTCCGGGTCAGTTCATGGTTATCTCTGGTGTTGGTGGTCTGGGTCACATCGCT
GTTCAGTACGCTGCTGCTATGGGTATGCGTGTTATCGCTGTTGACATCGCTGACGACAAACTGGAACTGGCTCGTAAACACGG
TGCTGAATTTACCGTTAACGCTCGTAACGAAGACCCGGGTGAAGCTGTTCAGAAATACACCAACGGTGGTGCTCACGGTGTTC
TGGTTACCGCTGTTCACGAAGCTGCTTTCGGTCAGGCTCTGGACATGGCTCGTCGTGCTGGTACCATCGTTTTCAACGGTCTG
CCGCCGGGTGAATTTCCGGCTTCTGTTTTCAACATCGTTTTCAAAGGTCTGACCATCCGTGGTTCTCTGGTTGGTACCCGTCAG
GACCTGGCTGAAGCTCTGGACTTCTTCGCTCGTGGTCTGATCAAACCGACCGTTTCTGAATGCTCTCTGGACGAAGTTAACGA
CGTTCTGGACCGTATGCGTAACGGTAAAATCGACGGTCGTGTTGCTATCCGTTACTAA

Engineered methanol dehydrogenase from *Bacillus methanolicus* (UniProt: I3DVX6, Q5L and an A363L modification) (Roth et al., 2019)

ATGCATCATCACCATCACCACACCAACACCCTGTCTGCTTTCTTCATGCCGTCTGTTAACCTGTTCGGTGCTGGTTCTGTTAAC
GAAGTTGGTACCCGTCTGGCTGACCTGGGTGTTAAAAAAGCTCTGCTGGTTACCGACGCTGGTCTGCACGGTCTGGGTCTGT
CTGAAAAAATCTCTTCTATCATCCGTGCTGCTGGTGTTGAAGTTTCTATCTTCCCGAAAGCTGAACCGAACCCGACCGACAAAA
ACGTTGCTGAAGGTCTGGAAGCTTACAACGCTGAAAACTGCGACTCTATCGTTACCCTGGGTGGTGGTTCTTCTCACGACGCT
GGTAAAGCTATCGCTCTGGTTGCTGCTAACGGTGGTAAAATCCACGACTACGAAGGTGTTGACGTTTCTAAAGAACCGATGGT
TCCGCTGATCGCTATCAACACCACCGCTGGTACCGGTTCTGAACTGACCAAATTCACCATCATCACCGACACCGAACGTAAAG
TTAAAATGGCTATCGTTGACAAACACGTTACCCCGACCCTGTCTATCAACGACCCGGAACTGATGGTTGGTATGCCGCCGTCT
CTGACCGCTGCTACCGGTCTGGACGCTCTGACCCACGCTATCGAAGCTTACGTTTCTACCGGTGCTACCCCGATCACCGACG
CTCTGGCTATCCAGGCTATCAAAATCATCTCTAAATACCTGCCGCGTGCTGTTGCTAACGGTAAAGACATCGAAGCTCGTGAAC
AGATGGCTTTCGCTCAGTCTCTGGCTGGTATGGCTTTCAACAACGCTGGTCTGGGTTACGTTCACGCTATCGCTCACCAGCTG
GGTGGTTTCTACAACTTCCCGCACGGTGTTTGCAACGCTGTTCTGCTGCCGTACGTTTGCCGTTTCAACCTGATCTCTAAAGTT
GAACGTTACGCTGAAATCGCTGCTTTCCTGGGTGAAAACGTTGACGGTCTGTCTACCTACGACGCTGCTGAAAAAGCTATCAA
AGCTATCGAACGTATGGCTAAAGACCTGAACATCCCGAAAGGTTTCAAAGAACTGGGTGCTAAAGAAGAAGACATCGAAACCC
TGGCTAAAAACGCTATGAAAGACCTGTGCGCTCTGACCAACCCGCGTAAACCGAAACTGGAAGAAGTTATCCAGATCATCAAA
AACGCTATGTAA

Engineered methanol dehydrogenase from *Cupriavidus necator N-1* version *CT4-1* (UniProt: F8GNE5)

ATGCATCATCACCATCACCACACCCACCTGAACATCGCTAACCGTGTTGACTCTTTCTTCATCCCGTGCGTTACCCTGTTCGGT
CCGGGTTGCGTTCGTGAAACCGGTGTTCGTGCTCGTTCTCTGGGTGCTCGTAAAGCTCTGATCGTTACCGACGCTGGTCTGCA
CAAAATGGGTCTGTCTGAAGTTGTTGCTGGTCACATCCGTGAAGCTGGTCTGCAAGCTGTTATCTTCCCGGGTGCTGAACCGA
ACCCGACCGACGTTAACGTTCACGACGGTGTTAAACTGTTCGAACGTGAAGAATGCGACTTCATCGTTTCTCTGGGTGGTGGT
TCTTCTCACGACTGCGCTAAAGGTATCGGTCTGGTTACCGCTGGTGGTGGTCACATCCGTGACTACGAAGGTATCGACAAATC
TACCGTTCCGATGACCCCGCTGATCTCTATCAACACCACCGCTGGTACCGCTGCTGAAATGACCCGTTTCTGCATCATCACCA
ACTCTTCTAACCACGTTAAAATGGTTATCGTTGACTGGCGTTGCACCCCGCTGATCGCTATCGACGACCCGTCTCTGATGGTTG
CTATGCCGCCGGCTCTGACCGCTGCTACCGGTATGGACGCTCTGACCCACGCTATCGAAGCTTACGTTTCTACCGCTGCTACC
CCGATCACCGACGCTTGCGCTGAAAAAGCTATCGTTCTGATCGCTGAATGGCTGCCGAAAGCTGTTGCTAACGGTGACTCTAT
GGAAGCTCGTGCTGCTATGTGCTACGCTCAGTACCTGGCTGGTATGGCTTTCAACAACGCTTCTCTGGGTTACGTTCACGCTA
TGGCTCACCAGCTGGGTGGTTTCTACAACCTGCCGCACGGTGTTTGCAACGCTATCCTGCTGCCGCACGTTTCTGAATTTAAC
CTGATCGCTGCTCCGGAACGTTACGCTCGTATCGCTGAACTGCTGGGTGAAAACATCGGTGGTCTGTCTGCTCACGACGCTG
CTAAAGCTGCTGTTTCTGCTATCCGTACCCTGTCTACCTCTATCGGTATCCCGGCTGGTCTGGCTGGTCTGGGTGTTAAAGCT
GACGACCACGAAGTTATGGCTTCTAACGCTCAGAAAGACGCTTGCATGCTGACCAACCCGCGTAAAGCTACCCTGGCTCAGGT
TATGGCTATCTTCGCTGCTGCTATGTAA
